# M1-derived extracellular vesicles polarize recipient macrophages into M2 and alter skeletal muscle homeostasis in a hyper-glucose environment

**DOI:** 10.1101/2023.10.03.560690

**Authors:** Stefano Tacconi, Francesco Vari, Carolina Sbarigia, Diana Vardanyan, Serena Longo, Francesco Mura, Federica Angilè, Audrey Jalabart, Daniele Vergara, Francesco Paolo Fanizzi, Marco Rossi, Elizabeth Errazuriz-Cerda, Christel Cassin, Rienk Nieuwland, Anna Maria Giudetti, Sophie Rome, Luciana Dini

**Affiliations:** CarMeN Laboratory (UMR INSERM 1060/INRA 1397), HCL, Lyon 1 University, Pierre-Bénite, FRANCE; Department of Biology and Biotechnology “C. Darwin”, Sapienza University of Roma, Roma, ITALY; Department of Biological and Environmental Sciences and Technologies (Di.S.Te.B.A.), University of Salento, Lecce, ITALY; Research Center of Nanotechnologies for Engineering (CNIS), Sapienza University of Roma, Roma, ITALY; Department of Basic and Applied Sciences for Engineering, University of Rome Sapienza, Roma, ITALY; Centre d’Imagerie Quantitative Lyon Est (CIQLE), Lyon 1 University, Lyon, FRANCE.; Laboratory of Experimental Clinical Chemistry, Department of Clinical Chemistry, and Amsterdam Vesicle Center, AMC, Amsterdam UMC, Amsterdam, NETHERLANDS

**Keywords:** extracellular vesicles, hyperglycemia, macrophage activation, lipid metabolism, skeletal muscle

## Abstract

**Background:** Macrophages release not only cytokines but also extracellular vesicles (EVs). EVs are small lipid-derived vesicles with virus-like properties transferring lipids, RNA and proteins between cells. Until now, the consequences of macrophage plasticity on the release and the composition of EVs have been poorly explored. In this study, we determined the impact of high-glucose (HG) concentrations on macrophage metabolism, and characterized their derived EV subpopulations. Finally, we determined whether HG-treated macrophage-derived EVs participate in immune responses and in metabolic alterations of skeletal muscle cells.

**Methods:** THP1-macrophages (M0) were treated with 15mM (MG15) or 30mM (MG30) glucose. M1 or M2 canonical markers, pro– and anti-inflammatory cytokines and lactate production were evaluated. Macrophage-derived EVs were characterized by TEM, flow cytometry, and 1H-Nuclear magnetic resonance spectroscopy for lipid composition. M0 macrophages or C2C12 muscle cells were used as recipients of MG15 and MG30-derived EVs. The lipid profiles of recipient cells were determined, as well as protein and mRNA levels of relevant genes for macrophage polarization or muscle metabolism.

**Results:** M0 released 2 populations of small and large EVs (sEVs, lEVs) with specific lipid profiles. Proportionally to the glucose concentration, glucose-treatment induced glycolysis in M0 macrophages which consequently shifted into a pro-inflammatory M1 phenotype, containing increased triacylglycerol and cholesterol content. Glucose also affected macrophage sphingolipid and phospholipid compositions. The lipid profile differences between sEVs and lEVs were abolished and represented the lipid profile alterations of MG15 and MG30 macrophages. Both sEVs and lEVs from M15 and M30 macrophages polarized M0 into anti-inflammatory M2, with increased contents of triacylglycerol and cholesterol. MG15 lEVs and sEVs induced insulin-induced AKT hyper-phosphorylation and accumulation of triacylglycerol in muscle cells, a state observed in pre-diabetes. Conversely, MG30 lEVs and sEVs induced insulin resistance in myotubes.

**Conclusions:** As inflammation involves first M1 macrophages, then the activation of M2 macrophages to attenuate inflammation, this study demonstrates that the dialog between macrophages through the EV route is an intrinsic part of the inflammatory response. In a hyperglycemic context, EV macrophages could participate in the development of muscle insulin-resistance and chronic inflammation.

## BACKGROUND

Extracellular vesicles (EVs) are lipid-derived vesicles released from cells that convey proteins, lipids, nucleic acids, and other metabolites among different cell types. All cells release EVs from the plasma membrane (PM) whereas eukaryotic cells also release EVs called ‘exosomes’ formed during the inward budding of the limiting membrane of the late endosomes called ‘multivesicular bodies’ (MVBs) [1]. Some subsets of MVBs can fuse with the plasma membrane and release their vesicular cargoes outside the cells. As the separation of these subtypes of EVs based on their composition is until now not definitively resolved, the International Society of Extracellular Vesicles (ISEV) recommends the terminology of large EVs (lEVs, >200nm) to consider EVs originated from PM, and small EVs (sEVs, <200nm) to consider EVs released from the endolysosomal pathway or small PM buddings [2, 3]. Until now, this nomenclature has permitted to identify subtypes of EVs with specific lipid, RNA, and protein compositions [4–7] confirming that this nomenclature is indeed able to discriminate EV subpopulations likely originating from different cellular compartment. Although poorly studied, data on adipocyte-derived EVs [4] or on skeletal muscle-derived EVs [8] indicate that lEVs and sEVs might have specific biological functions.

Interestingly, though immune cells are the largest purveyor of EVs in the blood, the respective functions of lEVs and sEVs released from immune cells are poorly understood. In addition, although immune cell-derived EVs are now recognized as important actors of the immune system [9, 10], the impact of the immune cell environment on the release of lEVs and sEVs is also unknown. Nevertheless, two studies demonstrated that in response to cytokines [11] or endotoxins [12] immune cells modulate their metabolism to secrete pro– or anti-inflammatory mediators, concomitantly paralleled by an increased release of EVs. Among immune cells, macrophages are key cells of innate immunity that participate also in tissue remodeling. Therefore macrophages have a pleiotropic role within tissues to maintain tissue homeostasis in health and diseases. To fulfill this role, they must adapt to their microenvironment which results in the production of different macrophage phenotypes. Until now the consequences of macrophage plasticity on the release and composition of EVs have been explored in the fields of cancer, infections, and tissue repair [13–17]. But of importance are the modifications of the macrophage nutritional environment. Indeed, lipid metabolism plays a major role in the activation of both M1 and M2 macrophages, *e.g*.; fatty acid oxidation is essential for the activation of inflammasome in M2 macrophages, and glycolysis fuels fatty acid oxidation in M1 macrophages [18]. Therefore metabolic alterations such as those obtained during the development of high-fat diet-induced obesity or associated with alteration of glucose homeostasis (*i.e*.; diabetes) are associated with the development of low-grade inflammation and macrophage infiltration and polarization inside metabolic tissues [19]. In that context, it was found that a high-glucose (HG) treatment induced the release of sEVs from macrophages (HG-sEVs). HG-sEVs induced activation and proliferation of mesangial cells, and secretion of extracellular matrix and inflammatory cytokines, compared to control-sEVs. HG-sEVs activated mesangial cells through the transfer of TGF-β1mRNA. This data suggests that a state of hyperglycemia may modify the release and the function of sEVs from macrophages which consequently participate in the progression of diabetic nephropathy [20]. The same group also demonstrated that HG-sEVs contained an increased concentration of miR-7002-5p which could induce dysfunction, autophagy inhibition, and inflammation in recipient mouse tubular epithelial cells or kidney [21]. Taken together, the before-mentioned studies have provided a proof-of-concept that glucose itself can modulate the release of extracellular vesicles from macrophages.

However, the functions of the lEVs and sEVs released from macrophages are unknown, and the consequences of the glucose treatment on macrophage metabolism underlying the modulation of EV release have not yet been determined. As EVs are mainly lipid-derived nanovesicles, we hypothesized that the modulation of EV release from macrophages in response to glucose may be associated with alterations in macrophage lipid metabolism. Therefore, in this paper, we have determined the impact of HG concentrations on macrophage phenotype and lipid metabolism. Then we studied how these lipidomic alterations can affect the lipid composition of macrophage sEVs and lEVs. Finally, we determined whether lEVs and sEVs could participate in the immune response and in metabolic alterations of skeletal muscle (SkM), a tissue known to be infiltrated by macrophages during hyperglycemic conditions [22].

## METHODS

### Cell culture conditions and treatments

Human THP-1 monocytes were amplified in RPMI-1640 (Cambrex, NJ) supplemented with 10% heat-inactivated Fetal Bovine Serum (FBS) (Cambrex, NJ), 2mM L-glutamine (Cambrex), 100IU/mL penicillin and streptomycin solution and 10.000U/mL amphotericin (antimycotic solution) (Sigma Aldrich, MA). When cells reached the concentration of 5×10^5^ cells/mL, they were differentiated into macrophages (M0) with 100ng/mL Phorbol 12-Myristate 13-Acetate (PMA) as previously reported [23]. Thereafter, M0 macrophages were treated with 15mM (MG15) or 30mM glucose (MG30) for additional 24h. As a positive control, PMA-differentiated THP1 cells were polarized in M1 macrophages with 100ng/mL lipopolysaccharides (LPS, Sigma Aldrich, MA) and 10ng/mL IFNγ (ThermoFisher Scientific, MA), and in M2 macrophages with 10ng/mL IL-4 for 24h. As a control of osmotic stress, cells were treated with 30mM mannitol.

Mouse myoblasts C2C12 cells (from ATCC® CRL-1772™) were routinely maintained in DMEM 4.5g/L glucose supplemented with 10% heat-inactivated FBS, 1000UI/mL penicillin, 1000UI/mL streptomycin, and 2mM L-glutamine (37°C, 5% CO_2_). At confluence, differentiation was induced with DMEM 4.5g/L glucose supplemented with 2% Horse Serum, for one week.

To assess the effects of lEVs and sEVs, on recipient cells, 2μg/mL of EVs were added to 1×10^6^ THP1-derived macrophages or C2C12 myotubes grown in EV-depleted medium. After EV treatment, myotubes were serum-starved for 4h in DMEM 1.5g/L glucose, treated with 100nM insulin (Sigma-Aldrich, MA) for 10min, and immediately lysed for protein extraction.

### EV isolation and sizing

For EV isolation, RPMI growth medium was centrifuged at 100.000g at 4°C overnight and filtered at 0.2μm to remove EVs from FBS. EVs extraction was realized by differential centrifugation: 500g (10min, RT), the resulting supernatant was centrifuged at 800g (10min, RT) and then at 2,000g (20min, RT) to remove cellular aggregates. To pellet lEVs, the resulting supernatant was centrifuged at 20,000g (20 min, 4°C). The supernatant was filtered through a 0.22μm filter (polyethersulfone membrane filter units, Thermo Fisher Scientific, MA) and then centrifuged at 100,000g (70min, 4°C) and rinsed in filtered phosphate buffer saline (PBS). The resulting pellet containing lEVs or sEVs was resuspended in 50μL PBS without calcium. As shown in additional file 1A, we validated by flow cytometry that sEV pellet was depleted of lipoproteins, a classical contaminant from the culture serum. EV proteins were quantified using the Bradford protein assay. A NanoSight (NTA, Malvern Instruments) was employed to measure particle size distribution in lEV and sEV fractions. The number of particles and their movements were recorded for 3×60sec and analyzed using the NS500 software. We also used flow cytometry to confirm vesicle size distribution in lEV and sEV fractions (see below).

### Cell viability assay

3-[4,5-dimethylthiazol-2-yl]-2,5 diphenyl tetrazolium bromide (MTT) assay was performed according to [24]. M0, MG15 and MG30 macrophages were incubated with 1 mg/mL MTT in RPMI-1640 (2h, 37°C and 5% CO2). Cells were then washed 3× in PBS (0.2M, pH 7.4) and the reduced MTT formazan crystals were solubilized with dimethyl sulfoxide (DMSO) (Carlo Erba, IT). Absorbance was measured at 570nm (Ultrospec 4000 UV/Visible Spectrophotometer, Pharmacia Biotech, IT).

### Scanning Electron Microscopy (SEM)

M0, MG15 and MG30 macrophages cultured on glass coverslips were fixed with 2.5% glutaraldehyde in 0.1 mol/L cacodylate buffer (pH 7.4, 1h, 4°C) and post-fixed with 1% OsO4 in the same buffer. Cells were dehydrated with acetone (25%, 50%, 70%, 90% and 100%) followed by Critical Point Dryer CPD EMITECH K850 (Quorum Technologies Ltd, UK). Stub-mounted specimens were gold-coated with a Balzers Union SCD 040 (Balzers Union, LI) and examined with a Zeiss EVO HD 15 scanning microscope (Ziess, DE).

### Transmission Electron Microscopy (TEM)

Cells were fixed and dehydrated with the same protocol used for SEM and embedded in Spurr resin. 60nm sections were examined under a Zeiss Auriga Scanning Electron Microscope (Zeiss, DE) equipped with the STEM module operating at 20kV.

After isolation, EVs were fixed with 0.1% of paraformaldehyde in PBS (30min, RT). Fixed EVs were stained with 2% uranyl acetate (7min, RT), loaded on formvar-carbon coated grids and observed with a Zeiss Auriga Scanning Electron Microscope (Zeiss, DE) equipped with the STEM module operating at 20 kV. Immunogold labelling to detect CD63 and CD81 at the surface of extracellular vesicles was realized as in [25].

### Western blot analysis

Cells were lysed RIPA buffer (NaCl 150mM, Tris-HCl 50mM pH 8, MgCl2 2mM, SDS 0.1%, Deoxycholic Acid 0.5%, NP40 1%) containing phenylmethylsulfonyl fluoride and protease inhibitor cocktail (ThermoFisher Scientific, MA). After 5min sonication, insoluble material was pelleted by centrifugation (10min, 13,000g, 4°C). Supernatants were transferred into a new tube, and protein concentration was measured with the Bradford method. 15-30μg of cell proteins, or 10μg EV proteins were separated by sodium dodecylsulfate-polyacrylamide gel electrophoresis (10-12% polyacrylamide, SureCast™ Acrylamide Solution, ThermoFisher Scientific, MA) and transferred onto PVDF membranes. The membranes were blocked with 5% nonfat dry milk and/or 3% bovine serum albumin (BSA) in Tris-buffered saline containing 0.1% Tween 20 (TTBS) (1h, RT). Then, membranes were incubated overnight at 4°C with primary antibodies (Additional Table 1A). After washing with TTBS, the membranes were incubated (1h, RT) with the horseradish peroxidase-conjugated secondary antibody diluted in TTBS in 5% nonfat dry milk and/or 3% BSA (Additional Table 1A). The immunoreactive bands were detected by using enhanced chemiluminescence (ECL) reagent (Immobilon Crescendo Western HRP substrate; Merck Millipore, DE). Band densities were quantified by densitometry using Image LabTM Version 6.0.1 2017 (Bio-Rad Laboratories, CA).

### Real-Time PCR

Total RNA extracted by using Trizol (Invitrogen, USA). 1.5μg of RNA was converted to cDNA using a single-step SuperScriptTM IV kit (Invitrogen, Carlsbad, CA) protocol. PCR was realized by using CFX ConnectTM Real-Time PCR Detection System (BIORAD, Hercules, CA). Primer sequences are in Additional Table 1B.

### Oil Red-O lipid droplet staining

Post EV treatment, cells were grown on 11mm glass coverslips, washed with DPBS, fixed 5min with 10% formalin, washed 5min with 60% isopropanol and left to completely dry. Subsequently, cells were incubated for 20min in 0.5% Oil Red-O/isopropyl alcohol and washed several times with DPBS. Then nuclei were stained with 1μM diamidino-2-phenylindole (DAPI). The coverslips were washed and visualized with a Zeiss Microscope equipped with Apotome (Zeiss, Germany). Images from different samples were recorded by using the same settings.

### Extracellular lactate measurement

The levels of lactate in the conditioned medium of M0, MG15, and MG30 macrophages were measured as reported in [26].

### Thin-Layer Chromatography (TLC)

Cell and EV total lipids were extracted by the Bligh and Dyer procedure. Lipids were loaded on silica gel plates for separation by thin-layer chromatography (TLC). Plates were developed with hexane/ethyl ether/acetic acid (70/30/1; v/v/v) for neutral lipids, with chloroform/methanol/water (65/25/4; v/v/v) for phospholipids and with toluene/methanol (70:30; v/v) for sphingolipids. After development, plates were uniformly sprayed with 10% cupric sulfate in 8% aqueous phosphoric acid, allowed to dry (10min, RT) and then placed into a 145 °C oven (10mn). The identification of different species was made by developing specific standards with the same experimental conditions. Spot intensity was measured by densitometry.

## ^1^H-Nuclear magnetic resonance (NMR) Spectroscopy

All measurements, on lipid extract from EVs, were performed on a Bruker Avance III 600 Ascend NMR spectrometer (Bruker, Biospin Milan, IT) operating at 600.13 MHz for 1H observation, equipped with a TCI cryoprobe incorporating a z-axis gradient coil and automatic tuning-matching. Experiments were acquired at 300 K in automation mode after loading individual samples on a Bruker Automatic Sample Changer, interfaced with the software IconNMR (Bruker). Lipid extracts were dissolved in 600 μL of CD_3_OD/CDCl_3_ (1:2 mix) and transferred into a 5mm NMR tube, using tetramethylsilane (TMS, δ=0.00ppm) as internal standard. A one-dimensional experiment (zg Bruker pulse program) was performed with 256 scans, 64K data points, a spectral width of 20.0276 ppm, 2s delay, p1 8 μs, and 2.73 acquisition time. The resulting FIDs were multiplied by an exponential weighting function corresponding to a line broadening of 0.3 Hz before Fourier transformation, automated phasing, and baseline correction. Lipid species were assigned based on 2D NMR spectra analysis (2D ^1^H Jres, ^1^H COSY, ^1^H-13C HSQC) and comparison with published data [27].

### Flow cytometry

Samples were pre-diluted in PBS to a count rate between 2,000 and 3,000 events/s to prevent swarm detection when triggering on the side scatter detector. Antibodies were diluted in PBS and antibody aggregates were removed by centrifugation at 18,890g, 20°C for 5 minutes before use. 20μL of pre-diluted samples were incubated with 2.5μL of lactadherin-FITC (Additional table 1A) for two hours at room temperature in the dark. The post-staining was diluted by adding 200 μL PBS to decrease background fluorescence from unbound reagents. All samples were analyzed by flow cytometer (A60-Micro, Apogee Flow Systems; UK) at a flow rate of 3.01μL/minute. Each sample was measured for four minutes or until 500,000 events were detected, triggering on 405nm side scatter using a threshold corresponding to a side scattering cross-section of 10nm2 (Rosetta Calibration; Exometry, NL). Flow cytometry scatter ratio (Flow-SR) was applied to determine the size and refractive index (RI) of particles and to improve specificity by enabling label-free differentiation between EVs and lipoprotein particles [28]. Data analyses were performed in Custom-build software (MATLAB R2018b, Mathworks, MA) and FlowJo (v10.6.1; FlowJo, OR).

### Statistical analyses

Data are expressed as Means ± SEM. Multiple comparisons were performed by two-way ANOVA. Comparisons between two groups were performed using a student’s *t*-test (GraphPad Prism 7 software, GraphPad Software, San Diego, CA). p<0.05 were considered significant.

## RESULTS

### Hyperglucose (HG) treatment induced macrophage polarization towards a pro-inflammatory phenotype and altered their lipid metabolism

To understand the effect of HG concentrations on the release of EVs from macrophages, we first characterized the phenotypes of HG-treated macrophages. Two high glucose concentrations were used, *i.e*.; 15mM (MG15 macrophages), to mimic a state of hyperglycemia, a risk factor for many diseases (*e.g*.; ischemic cardiovascular injuries [29], renal diseases [30], sepsis [31], cancer [32] and diabetes), and 30mM (MG30 macrophages), known to activate the NLRP3 inflammasome in M0 THP-1 macrophages, to mimic a state of a persistent inflammatory response (*e.g*.; in chronic wounds). Of note, we validated that the two glucose concentrations did not affect macrophage viability (Additional Fig. 1B). After glucose treatment M1, M2 markers, and inflammatory cytokines were quantified. As shown in Fig.1A-B, M0 THP-1 macrophages are composed of M1 and M2 macrophages which shifted to an M1-enriched population; *i.e*.; increased level of CD86 proportional to the glucose concentration (Fig. 1A). In agreement, these pro-inflammatory M1 had increased expressions of pro-inflammatory cytokines (*i.e*.; IL-1beta and IFNα, Fig. 1C) and no modulation of the immune-suppressive IL-10 cytokine (Fig. 1C). The concomitant increase in TLR4 and NFκB expressions further validate the polarization of M0 THP-1 into M1 macrophages (Fig. 1D-E, [33]). We validated that the HG effect on macrophage polarization was independent of osmotic stress as 30mM mannitol did not reproduce these data (Additional Fig. 1C).

**Figure 1:**
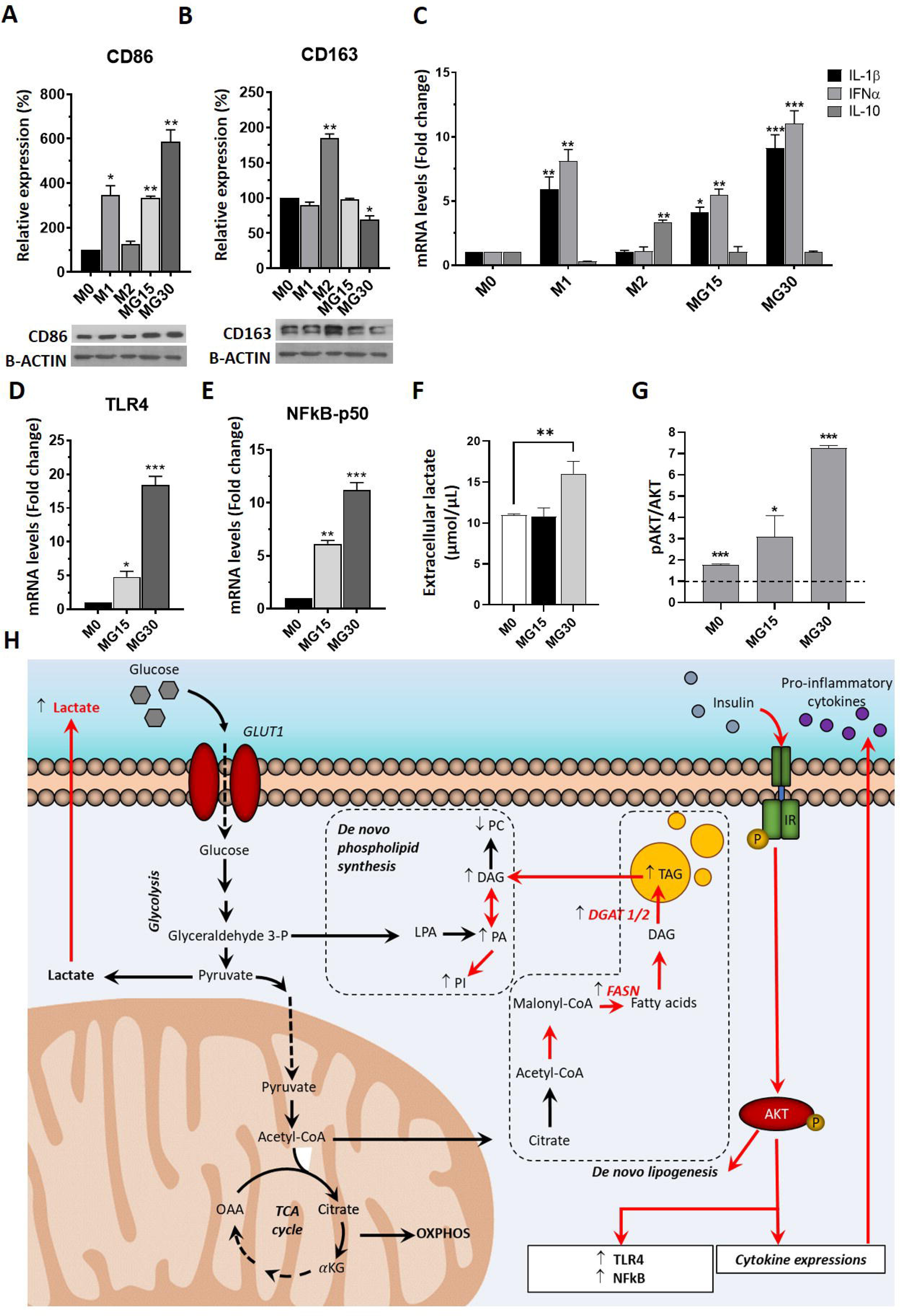
High glucose polarizes macrophages into M1. Protein levels of CD86 **(A)** and CD163 **(B)** determined by Western Blot in THP-1-derived macrophages. M0=untreated macrophages, MG15=15 mM glucose, MG30=30 mM glucose. Values are expressed as % of M0. (**C**) mRNA levels of IL-1β, Interferon α (IFNα) and IL-10, (**D**) toll-like receptor 4 (TLR4) and (**E**) subunit p-50-nuclear factor kappa-light-chain-enhancer of activated B cells (p50-NFkB) in macrophages. Data are normalized to glyceraldehyde 3-phosphate dehydrogenase (GAPDH) mRNA level, then expressed as fold changes of M0. M1=THP-1-derived macrophages treated with IFNγ and lipopolysaccharide (LPS) were used as a positive control of M1 polarization, M2=THP-1-derived macrophages treated with IL-4 used as a positive control of M2 polarization. **(F)** Lactate concentrations in the macrophage-conditioned medium. **(G)** Western Blot analysis of AKT phosphorylation in response to insulin stimulation expressed to basal pAKT. **(H)** Summary of the effect of glucose treatement on macrophage lipid metabolism. values are means ± SEM (n=3); *p* values are from student *t*-test (stimulated *vs* untreated), (∗) *p* < 0.05, (∗∗) *p* < 0.01, (∗∗∗) *p*< 0.001. GLUT1: glucose transporter 1; LPA: lysophosphatidic acid; PC: phosphatidylcholine; PA: phosphatidic acid; PI: phosphatidylinositol; DAG: diacylglycerol; TAG: triacylglycerol; FASN: fatty acid synthase; DGAT 1/2: diacylglycerol acyltransferase 1/2; IR: insulin receptor.

It is known that macrophage polarization is strongly associated with modifications of metabolism [34]. Therefore, to determine how HG might impact EV release, we determine how HG treatments affected macrophage metabolism. MG30 macrophages had an increased concentration of lactate in the conditioned medium (Fig. 1F), consistent with the fact that macrophage polarization into M1 is associated with a shift into a glycolytic metabolism [35]. Conversely, MG15 macrophages did not overproduce lactate suggesting a dose-dependent response of macrophages toward the glucose concentrations. HG did not affect the insulin sensitivity of MG15 and MG30 macrophages which had increased phosphorylation of insulin-induced AKT (Fig. 1G, Additional Fig. 1D). AKT is a key protein involved in the inhibition of mitochondrial oxidative phosphorylation [36] which is decreased in glycolytic M1 macrophages (Fig. 1H). Therefore these metabolic data supported the polarization of HG-treated macrophages into an M1 phenotype, with a stronger effect of 30mM *vs* 15mM glucose.

As it is known that glycolysis fuels the synthesis of fatty acids (FA) [37], we quantified the expressions of key proteins involved in lipid storage and synthesis in macrophages (Fig. 1H). As expected, MG15 and MG30 macrophages had increased expression of the FASN protein (Fig. 2A-B) confirming the endogenous synthesis of FA [38]. Concomitantly, DGAT2 protein was increased in MG15 and MG30 macrophages (Fig. 2A-C) and DGAT1 in MG30 (Fig. 2A-D) showing that macrophages accumulated lipids in response to HG, independent from the glucose concentration. Indeed, Oil-Red-O labelling (Fig. 2E) and lipid analyses (Fig. 2F, Additional Fig. 1E) demonstrated the huge accumulation of triacylglycerols (TAG) in HG-treated macrophages. Cells can also mobilize TAG to produce precursor FA that are used for phospholipid synthesis (Fig. 1H). In agreement, PA and PI are increased in MG15 and MG30 macrophages (Fig. 2F, Additional Fig. 1E).

**Figure 2:**
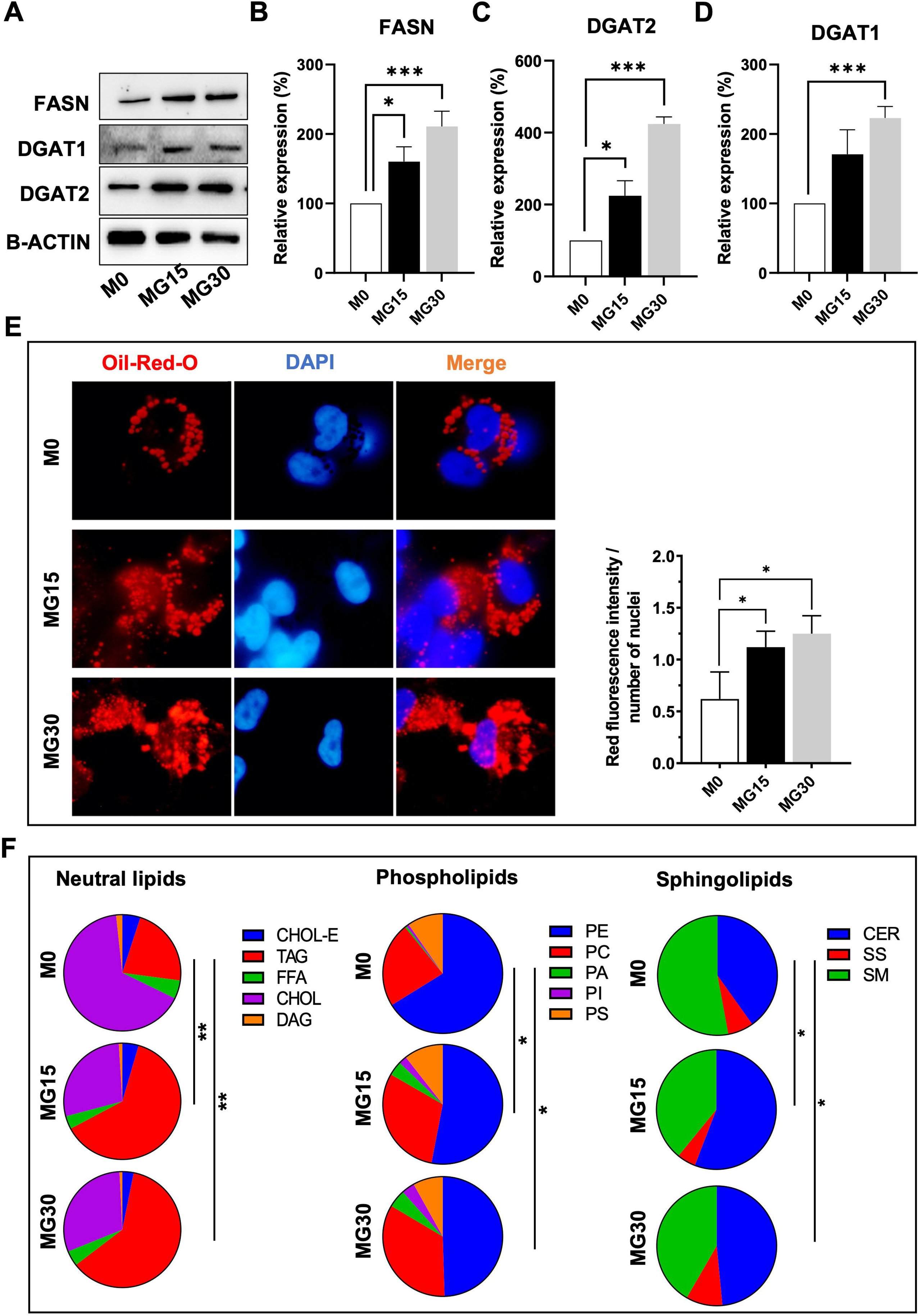
High glucose modifies macrophage lipid distribution. Western blot analysis (representative images of the blots in (**A**)) of fatty acid synthase (FASN) (**B**), diacylglycerol acyltransferase (DGAT) 2 (**C**) and DGAT1 (**D**) in M0, MG15 and MG30 macrophages. Values were normalized to beta-actin and reported as % of M0. Values are means ± SEM (n=3); *p* values are from student *t*-test (stimulated *vs* untreated), (∗) *p* < 0.05, (∗∗) *p* < 0.01, (∗∗∗) *p*< 0.001. M0=untreated macrophages, MG15=15 mM glucose, MG30=30 mM glucose. (**E**) Oil-red oil and 4′,6-diamidino-2-phenylindole (DAPI) staining of M0, MG15 and MG30 macrophages (**left**) and relative quantification of fluorescence intensity (**right**). (**F**) Distribution of neutral lipids, phospholipids, and sphingolipids in M0, MG15 and MG30 macrophages. Data are expressed as % and significantly different lipid distributions were identified with a chi-squared test. CHOL-E: cholesterol ester; TAG: triacylglycerol; FFA: free fatty acid; CHOL: cholesterol; DAG: diacylglycerol; PE: phosphatidylethanolamine; PC: phosphatidylcholine; PA: phosphatidic acid; PI: phosphatidylinositol; PS: phosphatidylserine; CER: ceramide; SS: sphingosine; SM: sphingomyelin.

HG also modulated the distribution of sphingolipids in macrophages. The ratio of sphingomyelin (SM)/ceramides (CER) was decreased in MG15 and MG30 compared to the untreated M0 macrophages (Fig. 2F, additional Fig. 1E). This accumulation of ceramides was in line with a previous study demonstrating that M1 macrophages expressed a ceramide-generation-metabolic pattern *vs* M2 macrophages [39]. Taken together, these data demonstrate that macrophages adapt their lipid metabolism in response to the increased extracellular concentrations of glucose, resulting in phospholipids remodeling and accumulation of ceramides.

### HG treatments affected the release and the lipid composition of macrophage-released EVs

As the HG treatment modifies the concentration of lipids involved in membrane synthesis and curvature [40], we suspected that this may affect the biogenesis and composition of macrophage-released EVs. SEM images revealed numerous buddings on the surface of macrophages (Fig. 3A). However, the plasma membrane of MG30 macrophages produced more buddings compared to MG15 or untreated M0 macrophages (Fig. 3B). As EVs were collected from the conditioned medium of HG-treated macrophages grown in EV-free medium, we validated that the EV-free medium did not affect macrophage viability (Additional Fig. 1F). Then LEVs and sEVs were purified from fresh conditioned medium (Fig. 3C left). Both lEVs and sEVs contained CD81 and CD63 (Fig. 3C right). We were not able to determine firmly the origin of the sEVs which could originate from small buddings in the plasma membrane or from late endosomal compartments. However, analysis of their lipid distribution by ^1^H-NMR indicated that they had a different origin from lEVs (Fig. 3D, Additional Fig. 2A, Additional table 2). Compared to lEVs, the sEVs were enriched in cholesterol, glycerophospholipids, TAG, and diacylglycerols (DAG), whereas lEVs were enriched in phospholipids and depleted in cholesterol compared to sEVs. The use of NTA to quantify particle sizes below 500nm, showed that M0 macrophages produced more lEVs than sEVs. In addition, lEVs were less homogeneous in particle size than sEVs (Fig. 3E, also validated by flow cytometry in Additional Fig. 1G).

**Figure 3.**
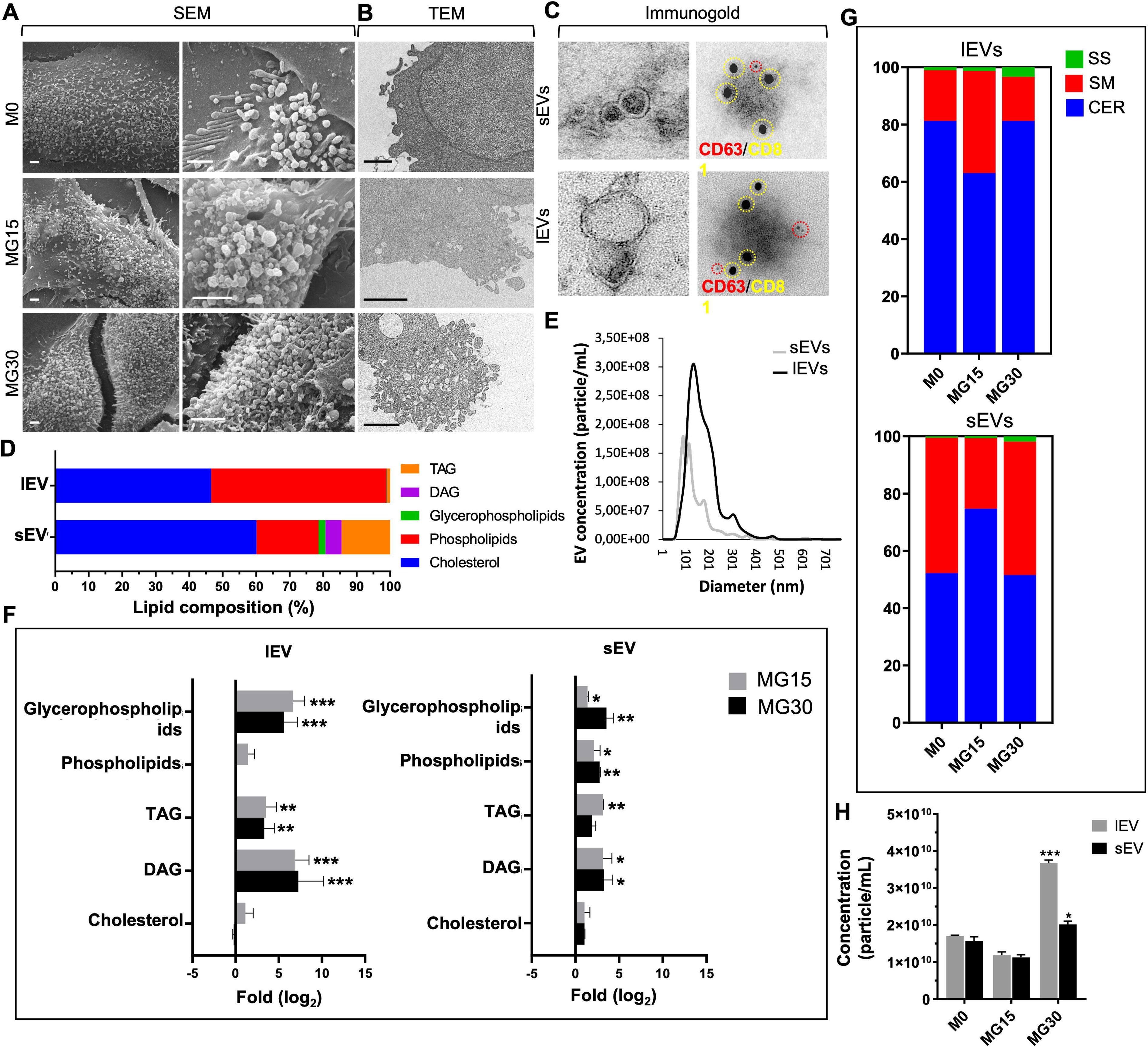
Macrophage-released EV, lipids, and biogenesis are altered by HG treatments. (**A**) Representative SEM images of the surface of macrophages. (**B)** Representative TEM images of the membranes of macrophages. In **A** and **B**, scale bar= 500 nm. (**C)** TEM images of lEVs and sEVs released from M0 macrophages (**left**). (**C**) **right**, immunogold labelling of macrophage-derived EVs to detect CD63 and CD81 at the EV surface. Red= CD63, 5nm gold particles, yellow= CD81, 15nm gold particles. (**D)** Lipid distribution in lEVs and sEVs determined by ^1^H-NMR Spectroscopy. DAG: diacylglycerol; TAG: triacylglycerol. (**E)** Size distribution of lEVs and sEVs determined by Nanotracking Analyses (NTA). (**F)** Lipid enrichment determined by^1^H-NMR, in lEVs and sEVs released from MG15 and MG30 macrophages compared to M0 macrophages. Values are expressed as log_2_ ratios of control M0. A representative ^1^H-NMR spectrum obtained at 600 MHz of CD_3_OD/CDCl_3_ lipid extracts of lEVs and sEVs is shown in Additional Fig.2 A. (**G)** Sphingolipid profile of lEVs and sEVs from M0, MG15 and MG30 macrophages performed by thin layer chromatography. Representative TLC runs are shown in Fig. Additional Fig.2 B. (**H**) NTA quantification of lEVs and sEVs, expressed as particle/ml, released from M0, MG15 and MG30 macrophages. Values are means ± SEM (n=3); *p* values are from student *t*-test (stimulated *vs* untreated), (∗) *p* < 0.05, (∗∗) *p* < 0.01, (∗∗∗) *p*< 0.001. M0= untreated macrophages, MG15=15 mM glucose, MG30=30 mM glucose.

Then we determined the effect of HG treatments on the lipid composition of macrophage-released EVs (Additional Fig. 2B). Fig. 3F presents the fold enrichment in each class of lipids *vs* lEVs and sEVs released from untreated M0 macrophages, respectively. Both lEVs and sEVs had significant enrichment in TAG/DAG and glycerophospholipids when released from MG15 and MG30 *vs* M0 macrophages (Fig. 3F). Therefore, the differential lipid enrichment lEVs and sEVs (Fig. 3D) was lesser pronounced after HG treatments. In cells, the biosynthesis of TAG/DAG and glycerophospholipids begins from the glyceraldehyde 3-P formed *via* glycolysis (Fig. 1H). Thus, the increase of these lipids in HG-treated macrophages-derived lEVs and sEVs reflected the metabolic shift in the producing macrophages. Sphingolipid analyses demonstrated that the SM/CER ratio was lower in lEVs than in sEVs when released from M0 macrophages (Fig. 3G). This differential distribution between the 2 types of EVs has been also observed for other cell types [4], and the quantity of ceramide in EVs seems a way to discriminate between lEVs and sEVs. After treatment with 15mM glucose, sEVs, and lEVs showed opposite modulations of their SM/CER ratios, *i.e*.; a decrease in lEVs (Fig. 3G) like in the releasing MG15 macrophages (Fig. 2E) but an increase in sEVs. For 30mM glucose, lEVs and sEVs had increased concentrations of sphingosine at the expense of SM, *vs* EVs from untreated M0 macrophages (Fig. 3G). Accumulation of sphingosine was also found in the releasing MG30 macrophages (Fig. 3G). Taken together, these data show that after HG treatment, the sphingolipid composition of lEVs reflected the sphingolipid composition of releasing macrophages. This was not the case for sEVs, further supporting that different mechanisms underlie the generation of lEVs and sEVs in macrophages. In addition, glucose treatment affected the release of EV from macrophages as for 30mM glucose, macrophages released more EVs *vs* untreated M0 or MG15 macrophages.

### LEVs and sEVs from HG-macrophages induced M2 polarization and regulated lipid metabolism of M0 recipient macrophages

*In vivo*, when released from cells, EVs are mainly captured by sentinel macrophages which constantly probe and clean their environment. Since HG-treated macrophages released a new population of EVs with altered lipid composition we hypothesized that it could have specific biological functions. Therefore, we treated M0 macrophages with EVs from HG-treated macrophages and analyzed the expressions of macrophage polarization markers and cytokines. lEVs and sEVs from MG30 or MG15 macrophages induced the expressions of CD163 (Fig.4A), IL-10 (Fig.4B), IL-6 (Fig. 4C) and decreased the expressions of TLR4 (Fig. 4D) and NFkB-p50 (Fig. 4E) in recipient M0 macrophages indicating that EVs from HG-treated M1 macrophages can polarize M0 recipient macrophages into M2 macrophages. In agreement, these recipient macrophages had increased protein levels of the main mitochondrial complexes, compared to untreated macrophages, confirming the metabolic shift toward an M2 phenotype (S3A).

**Figure 4.**
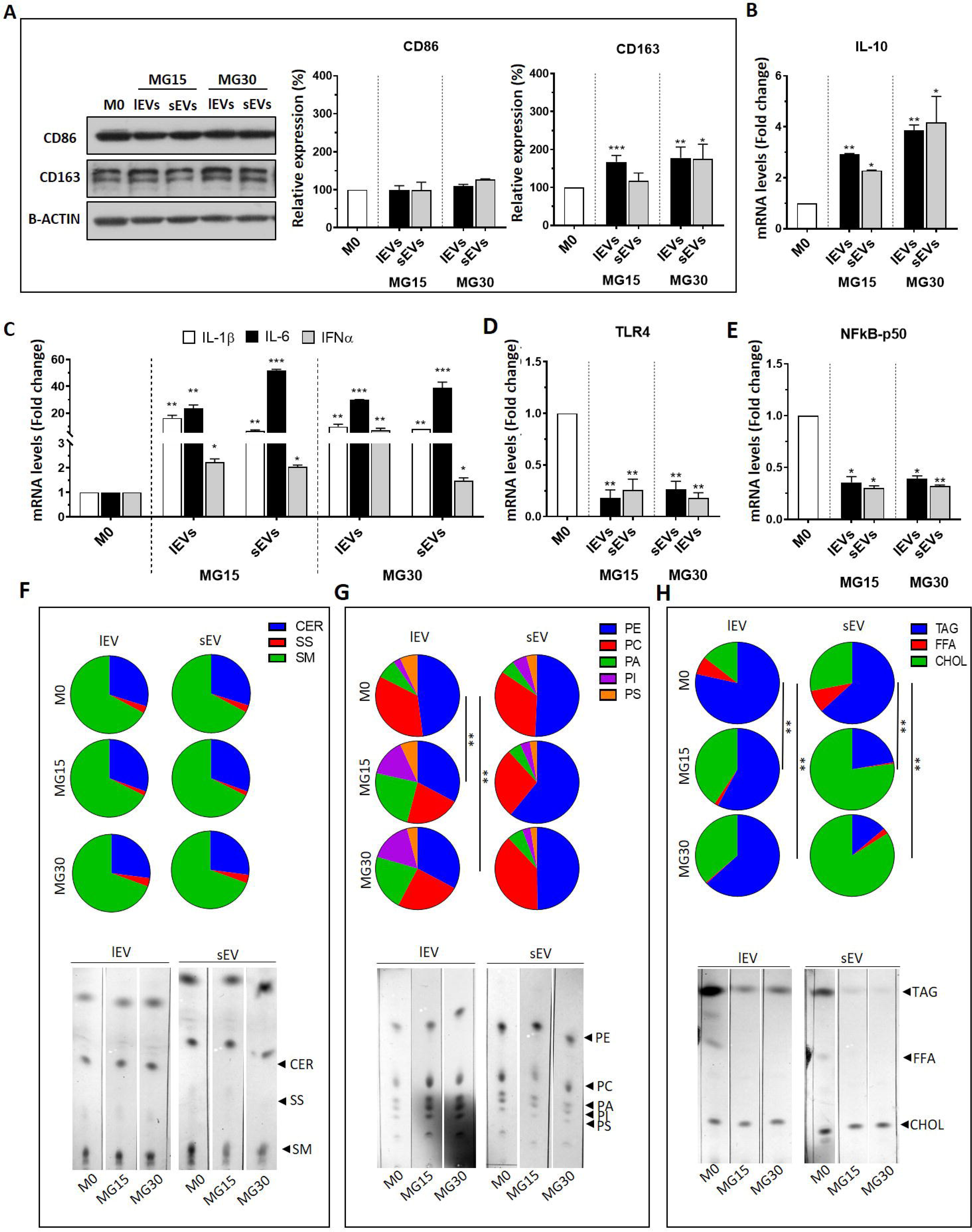
EVs from high glucose-treated macrophages affect M0 macrophage polarization and lipid composition. (**A**) Identification of M1 and M2 markers, CD86 and CD163 respectively, by Western blot. mRNA levels of interleukin (IL)-10 (**B**), IL-1β, IL-6, interferon α (IFNα) (**C**), toll-like receptor 4 (TLR4) (**D**), and subunit p-50-nuclear factor kappa-light-chain-enhancer of activated B cells (p50-NFkB) (**E**), in macrophages treated with lEVs and sEV released from M0, MG15 and MG30 macrophages. Data are normalized to glyceraldehyde 3-phosphate dehydrogenase (GAPDH) mRNA level, then expressed as fold changes of M0. Values are means ± SEM (n=3); *p* values are from student *t*-test (stimulated *vs* untreated), (∗) *p* < 0.05, (∗∗) *p* < 0.01, (∗∗∗) *p*< 0.001. (**F**) Profiling of neutral lipids, (**G**) phospholipids and (**H**) sphingolipids of EV-treated macrophages. Significantly different lipid distributions were identified by a chi-squared test. CER: ceramide; SS: sphingosine; SM: sphingomyelin; PE: phosphatidylethanolamine; PC: phosphatidylcholine; PA: phosphatidic acid; PI: phosphatidylinositol; PS: phosphatidylserine; TAG: triacylglycerol; FFA: free fatty acid; CHOL: cholesterol.

Then we analyzed the lipid profile of EV-treated macrophages to determine how EV treatment affected the lipid metabolism of the recipient macrophages. Neither sEVs nor lEVs from MG15 and MG30 macrophages modified the relative distribution of sphingolipids in recipient macrophages (Fig. 4F). This data indicated it is unlikely that sEVs or lEVs from MG15 and MG30 macrophages transfer sphingolipids (Fig.3G) in recipient macrophages. The treatment with lEV, but not sEV, leads to a decrease in the proportions of phosphatidylcholine (PC) and phosphatidylethanolamine (PE) in recipient M0 macrophages (Fig. 4G). As the decrease of both phospholipids is usually observed in activated macrophages [41], this result supports the effect of macrophage-derived EVs as actors of macrophage activation.

LEVs, but not sEVs, from MG15 and MG30 macrophages induced TAG accumulation (Fig. 4H) in recipient macrophages. This result could be related to the higher level of TAG/DAG in lEVs than sEVs, after HG-treatment (Fig. 3F). On the other side, sEVs from MG15 and MG30 macrophages induced much more accumulation of cholesterol inside recipient macrophages than lEVs (Fig.4H) also in agreement with the higher level of cholesterol in sEVs than lEVs (Fig. 3E-F). These data suggest that lEVs and sEVs could transfer TAG and cholesterol into recipient macrophages.

### EVs from HG-treated macrophages modulate metabolism and secretome of recipient muscle cells

As EVs from glucose-treated macrophages polarized M0 macrophages into an M2-like phenotype, we wondered whether they could also modulate the metabolism of non-immune cells. In this paper, we have considered skeletal muscle cells (SkM) as recipients. Indeed, in response to hyperglycemia, macrophages infiltrate SkM and release inflammatory cytokines, which can induce muscle wasting and impair muscle function [42]. Therefore, whether EVs from HG-treated macrophages can affect SkM homeostasis similarly to macrophage-released cytokines, is relevant. As shown in Fig. 5A (and additional Fig. 3B), both lEVs and sEVs from MG15 macrophages induced a hyperphosphorylation of AKT in response to insulin and accumulation of lipid droplets inside recipient SkM cells, already at 24h post-EV treatments (Fig. 5B). Conversely, recipient myotubes treated with lEVs and sEVs from MG30 macrophages strongly reduced insulin sensitivity (Fig. 5A) and lipid storage (Fig. 5B) demonstrating that they had a more pronounced effect on muscle energy balance and fat storage than lEVs and sEVs from MG15 macrophages.

**Figure 5:**
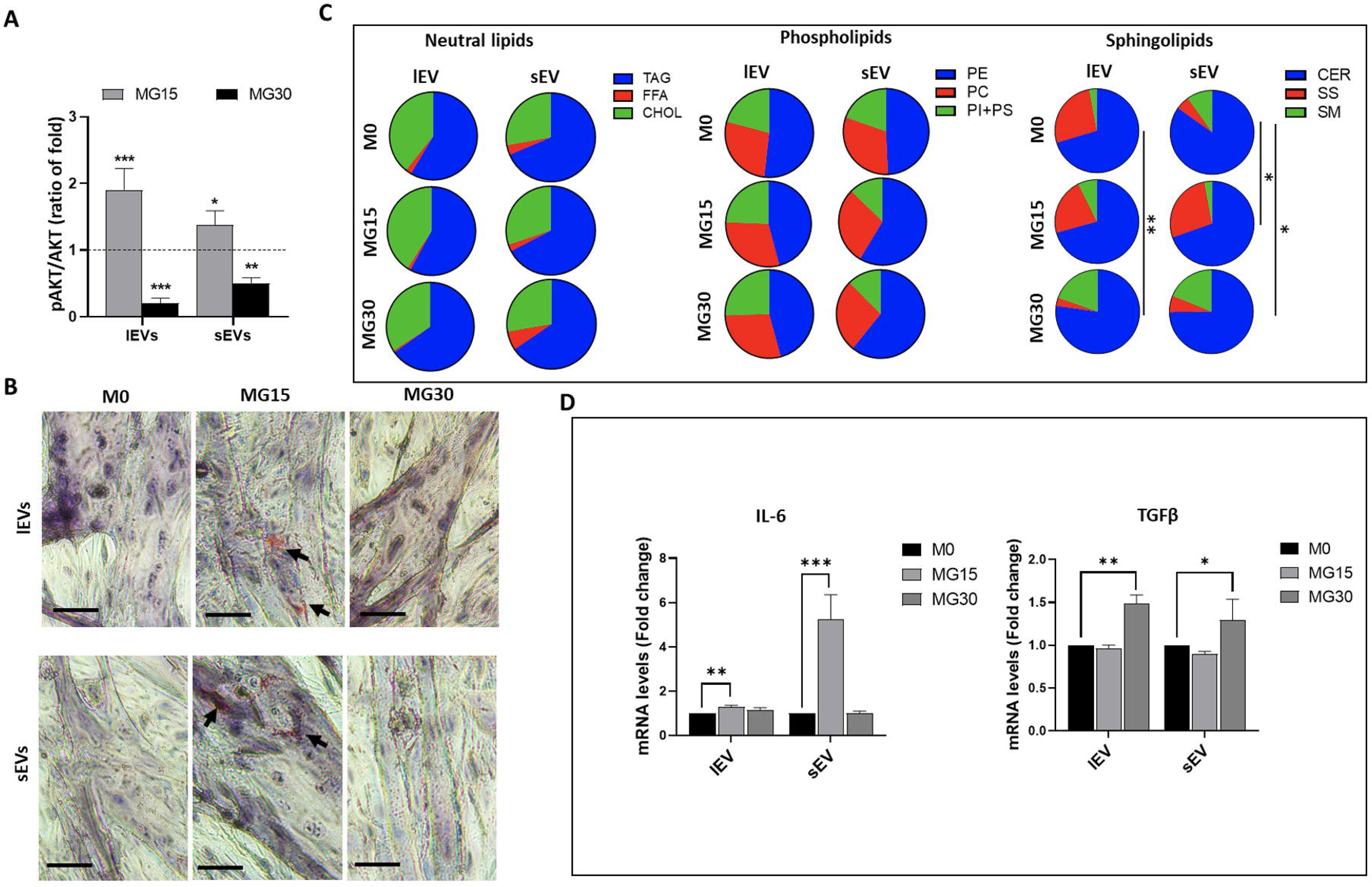
EVs from high glucose-treated macrophages modulate muscle homeostasis. (**A**) Quantification of insulin-induced phosphorylated AKT in C2C12 myotubes pre-treated with lEVs and sEVs from M0, MG15, or MG30 macrophages. Data are expressed as ratios (pAKT/AKT)_EV treated_/(pAKT/AKT)_untreated_. Images of the blot are shown in S3 B. **(B)** Lipid droplets detected by Oil-O-Red in C2C12 myotubes pre-treated with lEVs and sEVs from M0, MG15 and MG30 macrophages. Scale bar = 40 µm. **(C)** TLC analyses of neutral lipids, phospholipids, and sphingolipids of C2C12 myotubes treated with lEVs and sEVs from M0, MG15 and MG30 macrophages. Significantly different lipid distributions were identified by a chi-squared test. TAG: triacylglycerol; FFA: free fatty acid; CHOL: cholesterol; PE: phosphatidylethanolamine; PC: phosphatidylcholine; PI+PS: phosphatidylinositol+phosphatidylserine; CER: ceramide; SS: sphingosine; SM: sphingomyelin. (**D**) mRNA level of IL-6 and (**E**) of transforming growth factor β (TGFβ), in C2C12 myotubes treated with lEVs and sEVs from M0, MG15, and MG30 macrophages. Data are normalized to TBP mRNA level, then are expressed as fold of mRNA from C2C12 treated with lEVs or sEVs from M0 macrophages. Values are means ± SEM (n=3); *p* values are from student *t*-test (stimulated *vs* untreated), (∗) *p* < 0.05, (∗∗) *p* < 0.01, (∗∗∗) *p*< 0.001.

As macrophages treated with MG15-or MG30-derived EVs had altered lipid profiles (Fig. 3) we determined whether muscle treated with macrophage EVs had also altered lipid profiles. Conversely to recipient macrophages (Fig. 4), no significant modifications in phospholipids, neutral lipids, and cholesterol profiles were found in myotubes treated with MG15 or MG30 macrophage-derived EVs (Fig. 5C, Additional Fig. 3C). In contrast, lEVs and sEVs from MG30 macrophages induced an increase in SM content in recipient myotubes *vs* M0 or MG15 macrophage-derived lEVs and sEVs (Fig. 5C, Additional Fig. 3C). SM is a sphingolipid found in cell membranes, particularly enriched in the plasma membrane. It has been shown that SM interacts with the insulin receptor and thus can affect its activity [43]. If the accumulation of sphingomyelin is excessive, it can lead to the formation of large, rigid lipid rafts, which can hinder membrane protein mobility and signaling [44]. Therefore, the SM increase may partly explain the alteration of insulin sensitivity that we observed in myotubes treated with MG30 macrophage-derived LEVs and sEVs (Fig. 5A).

Then we determined whether MG15-or MG30-derived EVs modified the release of myokines from muscle cells. Both lEVs and sEVs from MG15 macrophages induced the release of IL-6 (Fig. 5D). In addition to its anti-inflammatory function IL-6 has also important regulatory roles by enhancing muscle insulin sensitivity [45]. This might explain why SkM cells treated with lEVs and sEVs from MG15 macrophages have increased insulin sensitivity (Fig. 5A). Conversely, both lEVs and sEVs from MG30 macrophages had a deleterious effect inducing the pro-fibrotic TGFβ expression in recipient SkM cells (Fig. 5E).

## DISCUSSION

Dysregulation of glucose homeostasis can result in elevated blood glucose levels, also known as hyperglycemia. It occurs in several metabolic and cardiovascular diseases, sepsis, and cancers and leads to oxidative stress, which in turn promotes inflammation by activating macrophages. Activated macrophages release pro-inflammatory cytokines that can lead to tissue damage and chronic inflammation. The effect of hyperglycemia on the differentiation and activation of subpopulations of macrophages has previously been investigated. It was found that HG-treatment on M1 macrophages pre-stimulated with LPS led to increased secretion of pro-inflammatory cytokines [46, 47], associated with an increase in ROS production and autophagy in macrophages. An HG environment also induced the secretion of pro-inflammatory cytokines during the differentiation of primary human macrophages [48]. In the present study, we show that differentiated naïve M0 macrophages grown in an HG environment expressed pro-inflammatory cytokines demonstrating the direct impact of a HG environment on macrophage polarization into M1.

In parallel, we found that glucose metabolism in M0 macrophages under HG conditions shifted towards a glycolytic metabolism, which had never been described before. This metabolic shift explains the polarization of M0 macrophages into M1, as unlike M2, M1 macrophages have anaerobic metabolism [49]. As M1 macrophages are also observed in situations of hypoxia (*e.g*.; in adipose tissue of obese patients [50] or during tumor initiation [51]), it was admitted that hypoxia was responsible for the induction M1 macrophages. The results of the present study demonstrate that an HG environment, without hypoxia, also induces a population of pro-inflammatory macrophages through the modulation of their glucose metabolism. Interestingly, the HG environment was not associated with the alteration of insulin-induced AKT phosphorylation in macrophages such as in metabolic tissues [52]. Consequently, in a diabetic hyperglycemic/hyperinsulinemic environment, AKT activity in macrophages may aggravate the *de novo* lipogenesis and TAG accumulation in macrophages [53].

This study demonstrated that M0 macrophages released a mixed population of sEVs and lEVs, based on their size distribution and morphologies, with very different lipid enrichment, suggesting specific mechanisms or plasma membrane budding locations for their genesis. Both lEVs and sEVs contained cholesterol, but sEVs are more enriched. As the technics used for lipid characterization in the producing cells and released EVs differ, we were not able to determine whether the differential enrichments in subspecies of phospholipids and sphingosines observed in HG-treated M0 macrophages are reflected on the EV phospholipids and sphingolipids compositions.

Our data show that HG-treated M0 macrophages released EVs enriched in glycerophospholipids and phospholipids, TAG, and DAG, and had a small increase in cholesterol. As EVs are mainly lipid-derived particles, this result indicates that locally at the site of inflammation, the M1 macrophage environment is enriched in these lipids. In the body, local accumulation of cholesterol (*i.e*.; the walls of blood vessels) and TAG (*i.e.*; adipose tissue) are signals for macrophage recruitment to the site of accumulation. Therefore, the release of lipid-derived EVs which disseminate lipids inside tissues, could be an additional mechanism together with cytokines in the development of chronic inflammation. Indeed, macrophages are phagocytic, and it has been demonstrated that exogenous EVs injected directly into the blood are cleared within two minutes by patrolling monocytes [54]. Previous studies have demonstrated that depending on the origin of the EVs, it can result in their polarization into M1 macrophages (*e.g*., adipocyte-derived EVs from diabetic subjects [55]; hepatocyte-derived EVs from NAFLD patients [56]; EVs from M1 macrophages induced by interferon-γ (IFN-γ) and LPS [57] or M2 (Human umbilical cord mesenchymal stem cell derived-EVs [58]; hypoxic glioma cell-derived EVs [59]). Here, we demonstrated that under HG conditions, M1 macrophages released EVs that can further polarize M0 recipient macrophages into M2 anti-inflammatory macrophages (*i.e*.; shift to oxidative metabolism and increase expressions of CD163 and of the anti-inflammatory IL-10). As the time-course of the inflammatory response involves first the activation of M1 macrophages in the early phase of inflammation, then the activation of M2 macrophages to attenuate inflammation and promote tissue repair, our data demonstrate that macrophage-derived EVs are an intrinsic part of the inflammatory response. In agreement with this study, it was recently demonstrated in a model of endometriosis in mice that M1 macrophage-derived EVs could repolarize macrophages into M2, inhibiting the development of endometriosis [60].

Hyperglycemia is common to many pathologies. Therefore, depending on the biological context the production of M2 macrophages from M1 macrophages through the EV route could be deleterious (*e.g*.; development of chronic inflammation or cancer) or could speed the process of tissue regeneration. In this latter case, it is known that after an injury early M1 macrophage infiltration participates in the clearance of necrotic debris, whereas M2 macrophage infiltration appears later to sustain tissue healing. Now, it is not known whether the two different macrophage populations result from a shift in macrophage polarization or from the recruitment of new monocytes. The data from this study and from [60] indicate that macrophage-derived EVs can shift macrophage phenotype.

Finally, we determined the effect of EV derived from macrophages on SkM cell homeostasis. MG15-derived EVs induced AKT hyperphosphorylation in response to insulin Excessive phosphorylation of AKT can result in its prolonged activation and insulin resistance, as cells become less and less responsive to the effects of insulin [61]. As activated AKT stimulates the synthesis of lipids, such as TAG, by promoting the uptake of fatty acids and their conversion into storage forms, it explains the accumulation of TAG in response to MG15-derived EVs vs untreated muscle cells. Conversely, EVs derived from macrophages treated with a higher concentration of glucose (30mM) completely altered insulin-induced AKT phosphorylation and TAG storage in recipient SkM cells suggesting that macrophage-derived EVs may participate in the development of insulin resistance associated with hyperglycemia [62]. In addition to muscle cell homeostasis, macrophage-derived EVs also modulate SkM secretome; *i.e*.; MG15-derived EVs induced ll-6 expression whereas MG30-derived EVs induced TGFβ in recipient muscle cells. IL-6 is produced by skeletal muscle cells as a result of stress, exercise, or injury. It has a protective effect on skeletal muscle cells (*e.g*.; involved in muscle regeneration or glucose uptake) but also acts as a signaling molecule between muscle and immune cells for muscle regeneration and adaptation [63]. Conversely, TGFβ has a deleterious effect on muscle cells as it induces fibrosis of skeletal muscle [64]. Therefore, these data demonstrate that EVs released by macrophages have an action on the muscle cell in relation to the glucose concentration used to treat the macrophages. For moderate hyperglycemia, macrophage-derived EVs could help to maintain muscle homeostasis, but for more severe hyperglycemia, they would participate in the development of muscle insulin resistance and alteration of muscle mass.

## CONCLUSIONS

This study demonstrates not only that a HG environment shapes macrophage phenotype by regulating their metabolism, but also the biological activity of the EVs released by these macrophages. Surprisingly, EVs derived from M1 in a HG environment induce M2 macrophages which can mitigate inflammatory response. Therefore in a context of chronic inflammation, these M1-derived EVs could have an unsuspected role in the development of skeletal muscle insulin-resistance associated with diabetes (Fig.6).

**Figure 6:** Graphical summary of the main results of this study

## Supplementary material

**Additional Figure 1**. (**A**) Flow cytometry quantification to detect lipoprotein contaminations (particles with RI>1.45) in the resuspended pellets containing lEV or sEV isolated from the conditioned medium of M0 macrophages. (**B**) MTT assay of THP-1-derived macrophages (M0) after the treatment with glucose 15 mM (MG15) or 30 mM (MG30). Values are expressed as % compared to untreated M0 cells. (**C**) Western Blot protein quantification of the polarization markers CD163 (M2) and CD86 (M1) in M0 macrophages treated with mannitol 30mM (osmolarity control) or with glucose 30mM. Values have been normalized with beta-actin and reported as % of M0. (**D**) Representative images of blots used for quantification of phosphorylated AKT (pAKT) in response to insulin, and total AKT in M0, MG15 and MG30 macrophages. (**E**) Representative TLC profile of neutral lipids, phospholipids, and sphingolipids in M0, MG15 and MG30 macrophages. (**F**) MTT assay of THP-1-derived macrophages (M0) treated for 24 h with EV-depleted medium (M0 + UC med) obtained by overnight ultracentrifugation (UC) at 110K xg. Values are expressed as % of untreated M0 macrophages. (**G**) Size distribution of lEVs and sEVs from M0 cells performed by dedicated flow cytometry. In **A, B, C, D, F** values are means ± SEM (n=3); *p* values are from student *t*-test (stimulated *vs* untreated), (∗) *p* < 0.05, (∗∗) *p* < 0.01, (∗∗∗) *p*< 0.001. CHOL-E: esterified cholesterol; TAG: triacylglycerol; FFA: free fatty acid; CHOL: cholesterol; DAG: diacylglycerol; PE: phosphatidylethanolamine; PC: phosphatidylcholine; PA: phosphatidic acid; PI: phosphatidylinositol; PS: phosphatidylserine; CER: ceramide; SS: sphingosine; SM: sphingomyelin.

**Additional Figure 2** (**A**) Representative 1H NMR spectra obtained at 600 MHz of CD3OD/CDCl3 lipid extracts of concentrated lEVs and sEVs. (**B**) Sphingolipids TLC analysis of lEVs and sEVs from M0, MG15 and MG30 macrophages. CER: ceramide; SS: sphingosine; SM: sphingomyelin.

**Additional Figure 3** (**A**) Western blots and quantification of complexes III, IV and V of the mitochondrial respiratory chain. Data were normalized to total protein levels (Ponceau S stain, on the bottom) and expressed as % of untreated M0 cells. values are means ± SEM (n=3); *p* values are from student *t*-test (stimulated *vs* untreated), (∗) *p* < 0.05, (∗∗) *p* < 0.01, (∗∗∗) *p*< 0.001. (**B**) Representative Blot images of Western blot analysis of pAKT/AKT with or without insulin stimulation in C2C12-derived myotubes treated with lEVs and sEVs from M0, MG15 and MG30 macrophages. (**C**) Representative TLC profile of neutral lipids, polar lipids and sphingolipids of C2C12 myotubes treated with lEVs and sEVs from M0, MG15 and MG30 macrophages. TAG: triacylglycerol; FFA: free fatty acid; CHOL: cholesterol; PE: phosphatidylethanolamine; PC: phosphatidylcholine; PI+PS: phosphatidylinositol+phosphatidylserine; CER: ceramide; SS: sphingosine; SM: sphingomyelin.

**Additional Table 1** (**A**) List of antibodies and probes used for Western Blot and Flow cytometry. (**B**) List of primers used for Real-Time PCR. H: human; M: mouse; R: rat.

**Additional Table 2** Chemical shift and assignments of main peaks in the ^1^H-NMR spectra of CDCl_3_/CD_3_OD extracts of sEVs and lEVs

## AUTHOR CONTRIBUTION

Conceptualization: ST, SR, LD; methodology and experiments: ST, AMG, FV, CS, DV, SL; fluorescence microscopy: ST; TEM analysis: EE, CC; SR, FM; SEM analysis: ST, FM; ^1^HNMR spectroscopy analysis: FA, FPF; Real Time PCR: ST, CS, AJ; data curation: ST, AMG, SR; writing original draft: ST, AMG, SR; writing/editing: DV, MR, RN, SR, LD.; supervision, SR, LD.

All authors have read and approved the submitted version of the manuscript.

## Data Availability Statement

All data from this article are included in this paper.

## Supporting information

Supplementary Images and tables

## ABBREVIATIONS

^1^H-NMR: ^1^H-nuclear magnetic resonance
AKT: protein kinase B
CD: cluster of differentiation
CER: ceramide
CHOL-E: cholesterol ester
CHOL: cholesterol
DAG: diacylglycerol
DGAT 1/2: diacylglycerol acyltransferase1/2
ECM: extracellular matrix
EVs: extracellular vesicles
FA: fatty acids
FASN: fatty acid synthase
FBS: fetal bovine serum
FFA: free fatty acid
GLUT1: glucose transporter 1
HG: high-glucose
IFN: interferon
IL: interleukin
ISEV: international society of extracellular vesicles
lEVs: large extracellular vesicles
LPA: lysophosphatidic acid
LPS: lipopolysaccharide
MTT: 3-[4,5-dimethylthiazol-2-yl]-2,5 diphenyl tetrazolium bromide
NTA: nano-tracking analysis
p50-NFkB: subunit p50 of the nuclear factor kappa-light-chain-enhancer of activated B cells
PA: phosphatidic acid
PBS: phosphate saline buffer
PC: phosphatidylcholine
PE: phosphatidylethanolamine
PI: phosphatidylinositol
PM: plasma membrane
PMA: phorbol 12-myristate 13-acetate
PS: phosphatidylserine
ROS: reactive oxygen species
RPMI: Roswell Park Memorial Institute medium
RT: room temperature
sEVs: small extracellular vesicles
SkM: skeletal muscle
SM: sphingomyelin
SS: sphingosine
TAG: triacylglycerol
TLC: thin layer chromatography
TLR4: toll-like receptor 4

## Acknowledgments

Sophie Rome and Rienk Nieuwland have trained Stefano Tacconi in the fields of EVs & metabolism and EVs & flow cytrometry, respectively.

