## Supplementary Images and tables for "M1-derived extracellular vesicles polarize recipient macrophages into M2 and alter skeletal muscle homeostasis in a hyper-glucose environment"

**Additional Figure 1**

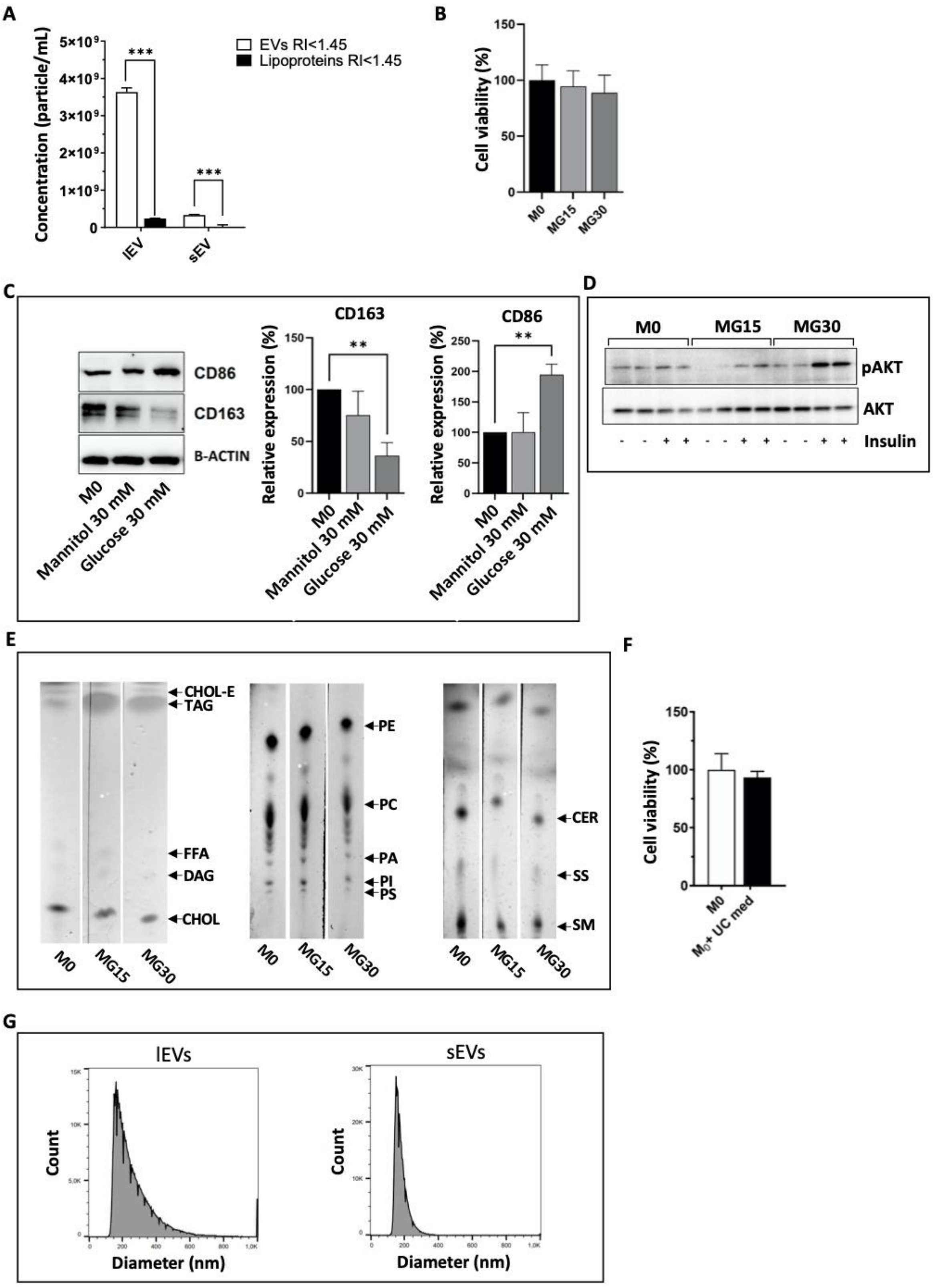

**Additional Figure 2**

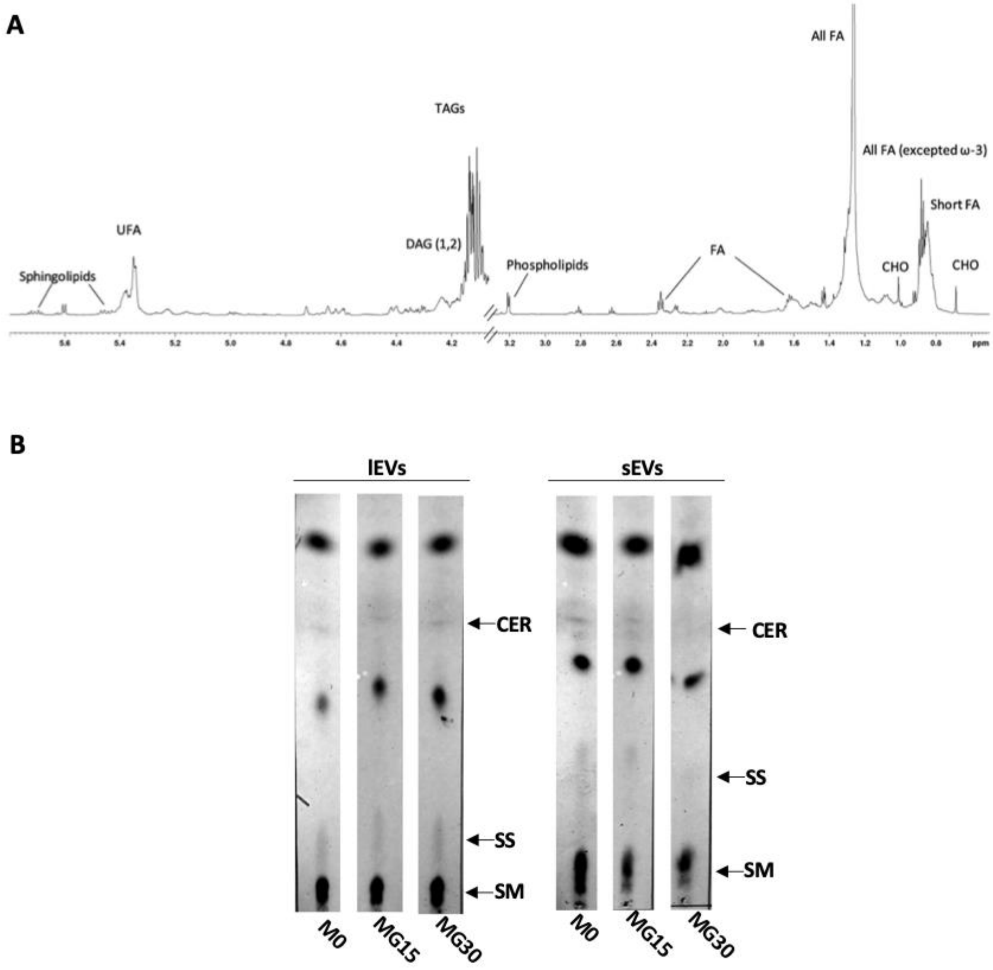

### Additional Figure 3

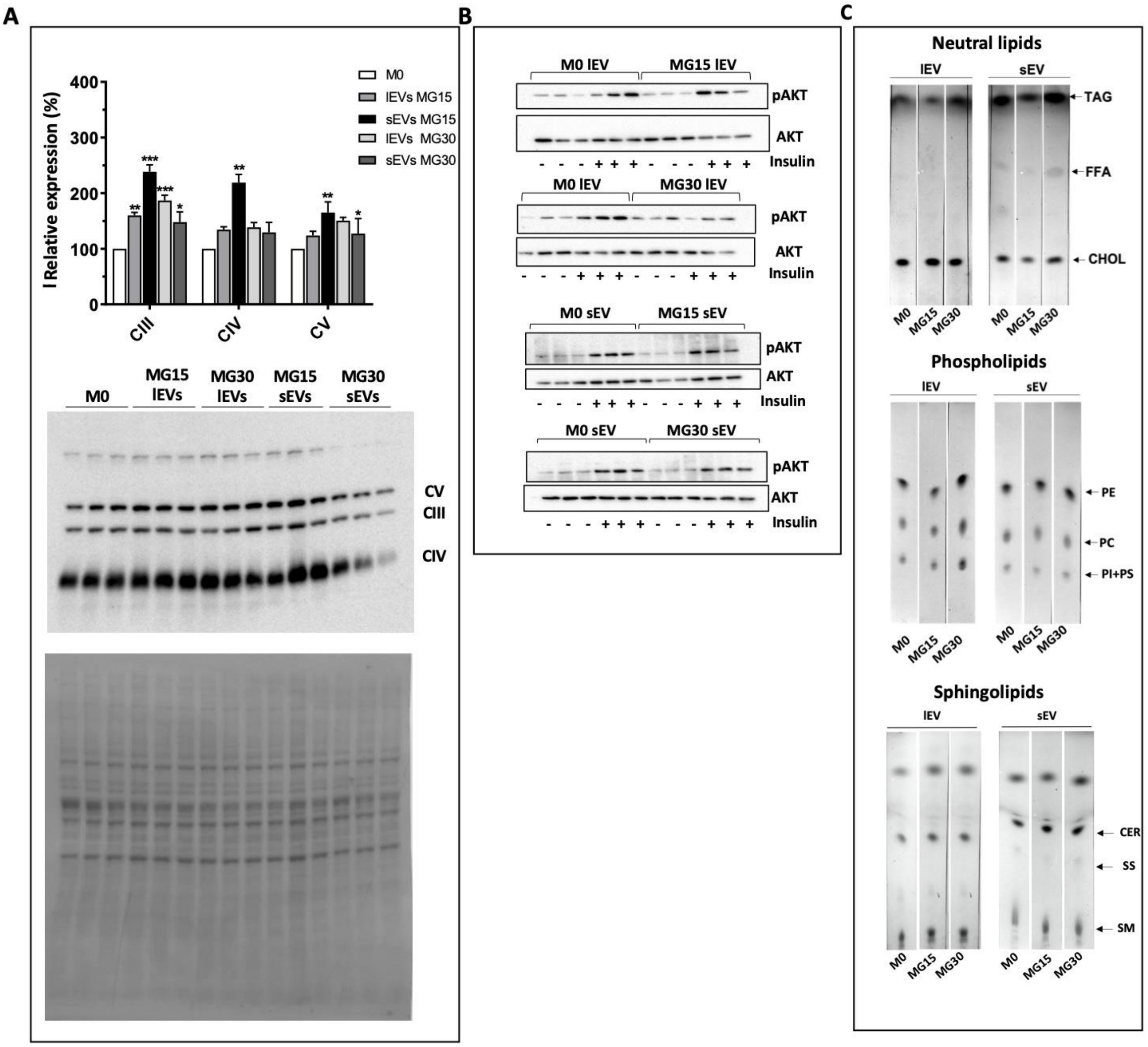

### Additional Table 1

**A**

| Antibody | Reactivity | Diluition | Reference code | Suppliers |
|---|---|---|---|---|
| anti-CD163 | H | 1:1000 | PA5-109327 | ThermoFisher Scientific, MA |
| anti-CD86 | H, M | 1:1000 | MA5-15697 | ThermoFisher Scientific, MA |
| Anti-AKT1 (phospho S473) | H, M, R | 1:1000 | ab81283 | Abcam, MA |
| Anti-pan-AKT | H, M, R | 1:1000 | ab8805 | Abcam, MA |
| Anti-AMPKα (phospho Thr172) | H, M, R | 1:1000 | 2531 | Cell Signaling Technology, MA |
| Anti-AMPKα | H, M, R | 1:1000 | 2532 | Cell Signaling Technology, MA |
| Anti-phospho-PDK1 (Ser241) | H, M, R | 1:1000 | 3061 | Cell Signaling Technology, MA |
| anti-PDK1 | H, M, R | 1:1000 | 3062 | Cell Signaling Technology, MA |
| anti-FASN | H, M, R | 1:1000 | ab128870 | Abcam, MA |
| anti-DGAT1 | H | 1:1000 | ab178711 | Abcam, MA |
| anti-DGAT2 | H, M | 1:1000 | ab237613 | Abcam, MA |
| OXPHOS Antibody Cocktail | H, M, R | 1:1000 | ab110413 | Abcam, MA |
| anti-β-actin | H, M, R | 1:2000 | A2228 | Sigma-Aldrich, St. Louis, MO |
| Anti-rabbit IgG, HRP-linked | R | 1:10000 | 7074 | Cell Signaling Technology, MA |
| Anti-mouse IgG, HRP-linked | M | 1:10000 | 7076 | Cell Signaling Technology, MA |
| **Probes** | | | | |
| Fitc-labelled Lactadherin (Flow cytometry) | | | NC0691117 | Haematologic Technologies, VT |

**B**

| Protein name | Target gene | Encoded protein | Forward | Reverse |
|---|---|---|---|---|
| Interleukin -1 beta | *IL1B* | IL-1β | 5′-GGCAATGAGGATGACTTGTT-3′ | 5′-TGTAGTGGTGGTCGGAGATT-3′ |
| Interleukin 10 | *IL10* | IL-10 | 5′-CGTGGAGCAGGTGAAGAATG-3′ | 5′-AGATCCGATTTTGGAGACCT-3 |
| Interferon alpha-1 | *IFNA1* | IFNα | 5′-GTGAGGAAATACTTCCAAAGAATCAC-3′ | 5′- TCTCATGATTTCTGCTCTGACAA-3′ |
| Nuclear factor kappa B subunit 1 | *NFKB1* | NFkB-p50 | 5’- GCACCTAGCTGCCAAAGAAG-3’ | 5’- CATGGCAGGCTATTGCTCATC-3’ |
| Toll-like receptor 4 | *TLR4* | TLR4 | 5’- CAACCAAGAACCTGGACCTG-3’ | 5’- AGGTAGAGAGGTGGCTTAGG-3’ |
| Glyceraldehyde-3-Phosphate Dehydrogenase | *GAPDH* | GAPDH | 5′-CTGCACCACCAACTGCTTAG-3′ | 5′-CTTCTGGGTGGCAGTGATGG-3′ |
| TATA-Box Binding Protein | *TBP* | TBP | 5’-TGGTGTGCACAGGAGCCAAG-3’ | 5’-TTCACATCACAGCTCCCCAC-3’ |

**Additional Table 2**

| Lipids | Assigment | $^1H$ ppm |
|---|---|---|
| **Cholesterol total** | $CH_3$-18<br>$CH_3$-21<br>$CH_3$-19 | 0.68<br>0.92<br>1.01 |
| **Phospholipids** | $-N(CH_3)_3$ | 3.19-3.21 |
| **TAG** | $-CH_2-$ | 4.17<br>4.31 |
| **DAG (1,2)** | $-CH_2-$ | 4.18<br>4.35 |
| **Glycerophospholipids** | Glycerol backbone sn1,3 | 4.40 |
